# Cannabis microbiome sequencing reveals several mycotoxic fungi native to dispensary grade cannabis flowers

**DOI:** 10.1101/030775

**Authors:** Kevin McKernan, Jessica Spangler, Lei Zhang, Vasisht Tadigotla, Yvonne Helbert, Theodore Foss, Douglas Smith

**Affiliations:** Medicinal Genomics Corporation, Woburn MA, 01801, USA

## Abstract

The Center for Disease Control estimates 128,000 people in the U.S. are hospitalized annually due to food borne illnesses. This has created a demand for food safety testing targeting the detection of pathogenic mold and bacteria on agricultural products. This risk extends to medical Cannabis and is of particular concern with inhaled, vaporized and even concentrated Cannabis products. As a result, third party microbial testing has become a regulatory requirement in the medical and recreational Cannabis markets, yet knowledge of the Cannabis microbiome is limited. Here we describe the first next generation sequencing survey of the microbial communities found in dispensary based Cannabis flowers and demonstrate the limitations in the culture based regulations that are being superimposed from the food industry.

## Introduction

Many states in the U.S. are crafting regulations for microbial detection on cannabis in absence of any comprehensive survey of Cannabis microbiomes. A few of these regulations are inducing growers to “heat kill” or pasteurize *Cannabis* flowers to lower microbial content. While this is a harmless suggestion, we must remain aware of how these drying techniques often create false negatives in culture based safety tests used to monitor colony-forming units (CFU). Even though pasteurization may be effective at sterilizing some of the microbial content, it does not eliminate various pathogenic toxins or spores. Aspergillus spores and mycotoxins are known to resist pasteurization ^1,2^. Similar thermal resistance has been reported for *E.coli* produced Shiga Toxin.^3^ While pasteurization may reduce CFU’s used in petri-dish or plating based safety tests, it does not reduce the microbial toxins, spores or DNA encoding these toxins.

Mycotoxin monitoring in Cannabis preparations is important since aflatoxin produced by Aspergillus species is a carcinogen. The clearance of aflatoxin requires the human liver enzyme CYP3A4 and this liver enzyme is potently inhibited by cannabinoids^4, 5^. Modern day cannabis flowers can produce up to 25% (w/v) cannabinoids presenting potent inhibition of CYP3A4 and CYP2C19. Health compromised patients exposed to aflatoxin and clearance-inhibiting cannabinoids raise new questions in regards to the current safety tolerances to aflatoxin. Similarly, Fusarium species are known to produce fungal toxins and has proven to be difficult to selectively culture with tailored media^6-8^. This is a common fault of culture-based systems as carbon sources are not exclusive to certain microbes and only 1% of microbial species are believed to be culturable^9^.

While these risks have been well studied in the food markets, the presence of the microbial populations present on cannabis flowers has never been surveyed with next generation sequencing techniques ^10-15^. With the publication of the Cannabis genome^16, 17^ and many other pathogenic microbial genomes, quantitative PCR assays have been developed that can accurately quantify fungal DNA present in Cannabis samples ^18^. Here, we analyze the yeast and mold species present in 10 real world, dispensary-derived Cannabis samples by quantitative PCR and sequencing, and demonstrate the presence of several mycotoxin producing fungal strains that are not detected by widely used culture-based assays.

## Results

A commercially available Total Yeast and Mold qPCR assay (TYM-PathogINDICAtor, Medicinal Genomics, Woburn) was used to screen for fungal DNA in a background of host *Cannabis* DNA. The TYM qPCR assay targets the 18S rDNA ITS (Internal Transcribed Spacer) region using modified primers described previously^19, 20^. Fungal DNA amplified using these primers may also be subjected to next generation sequencing to identify the contributing yeast and mold species. ITS sequencing has been widely used to identify and enumerate fungal species present in a given sample^21^.

We purified DNA from Cannabis samples obtained from two different geographic regions (Amsterdam and Massachusetts) several years apart (2011 and 2015). The majority of samples purified and screened with ITS qPCR were negative for amplification signal implying reagents clean of fungal contamination. Six of the 17 dispensary-derived Cannabis samples tested positive for yeast and mold in the TYM qPCR assay. These results were compared with the results derived from three commercially available culture based detection systems for each of the 17 samples (3M Petrifilm™ 3M Microbiology, St. Paul, MN, USA, SimPlates™ Biocontrol Systems, Bellevue, WA, USA, BioLumix™ Neogen, Lansing MI, USA). Of the 6 qPCR positive samples, two tested negative in all 3 culture-based assays and four tested negative in 1 or 2 of the culture-based assays (Table. 1). None of the qPCR negative samples tested positive in any of the culture based assays. Each of the 6 discordant samples was subjected to ITS sequencing to precisely identify the collection of microbes present. Four additional samples from a different geographic origin (Amsterdam) were also subjected to ITS sequencing, for a total of 10 Cannabis samples.

**Table 1.** Samples were cultured with 3 different techniques and compared to quantitative PCR (qPCR). Biolumix had the lower sensitivity failing to pick up 4/17 samples detected with other culture-based platforms. qPCR identified 2 samples that were not picked by any other method. Positive qPCR samples were sequenced to identify the contributing signal. Highlighted samples fail the 10,000 CFU/g cutoffs which equates to a Cq of 26 on the qPCR assay according to the manufacturers instructions. (f) is Fail or over 10,000 CFU/g. (p) is Pass or under 10,000 CFU/g.

| Samples | Yeast/Mold |  |  |  |
| --- | --- | --- | --- | --- |
|  | Simplate®(CFU/g) | 3M™ (CFU/g) | Biolumix | Cq |
| KD4 | 0 | 0 | pass | 21.71 (f) |
| KD8 | 0 | 0 | pass | 22.5 (f) |
| PC3 | 0 | 0 | pass | >40 (p) |
| White Widow | 0 | 0 | pass | >40 (p) |
| KD1 | 0 | 0 | pass | 29.33 (p) |
| KD2 | 0 | 0 | pass | >40 (p) |
| KD3 | 0 | 0 | pass | 30.16 (p) |
| KD5 | 1000 (p) | 6000 (p) | pass | 27.76 (p) |
| KD6 | 3000 (p) | 19000 (f) | pass | 24.72 (f) |
| KD7 | 0 | 0 | pass | >40 (p) |
| Liberty Haze | 172000 (f) | 89000 (f) | pass | 24.02 (f) |
| Blueberry Kush | 0 | 0 | pass | 37.99 (p) |
| Blueberry Kush<br>(Experimental) | >738000 (f) | too high (f) | fail | 15.71 (f) |
| Girl Scout Cookies | >738000 (f) | Too high (f) | pass | 19.66 (f) |
| Jake's Grape | >738000 (f) | Too high (f) | pass | 24.56 (f) |
| Serious Happiness | 0 | 0 | pass | >40 (p) |
| White Rhino | 0 | 3000 (p) | pass | >40 (p) |

Each discordant sample presented with an array of microbial species, as shown in figure 2. No sample presented with a single dominant species, and each sample displayed multiple species of interest. Of particular concern were the identified DNA sequences from toxin producing species: *Aspergillus versicolor*^22-26^, *Aspergillus terreus*^27^, *Penicillium citrinum*^23-30^, *Penicillium paxilli*^31,32^.

**Figure 1A.**
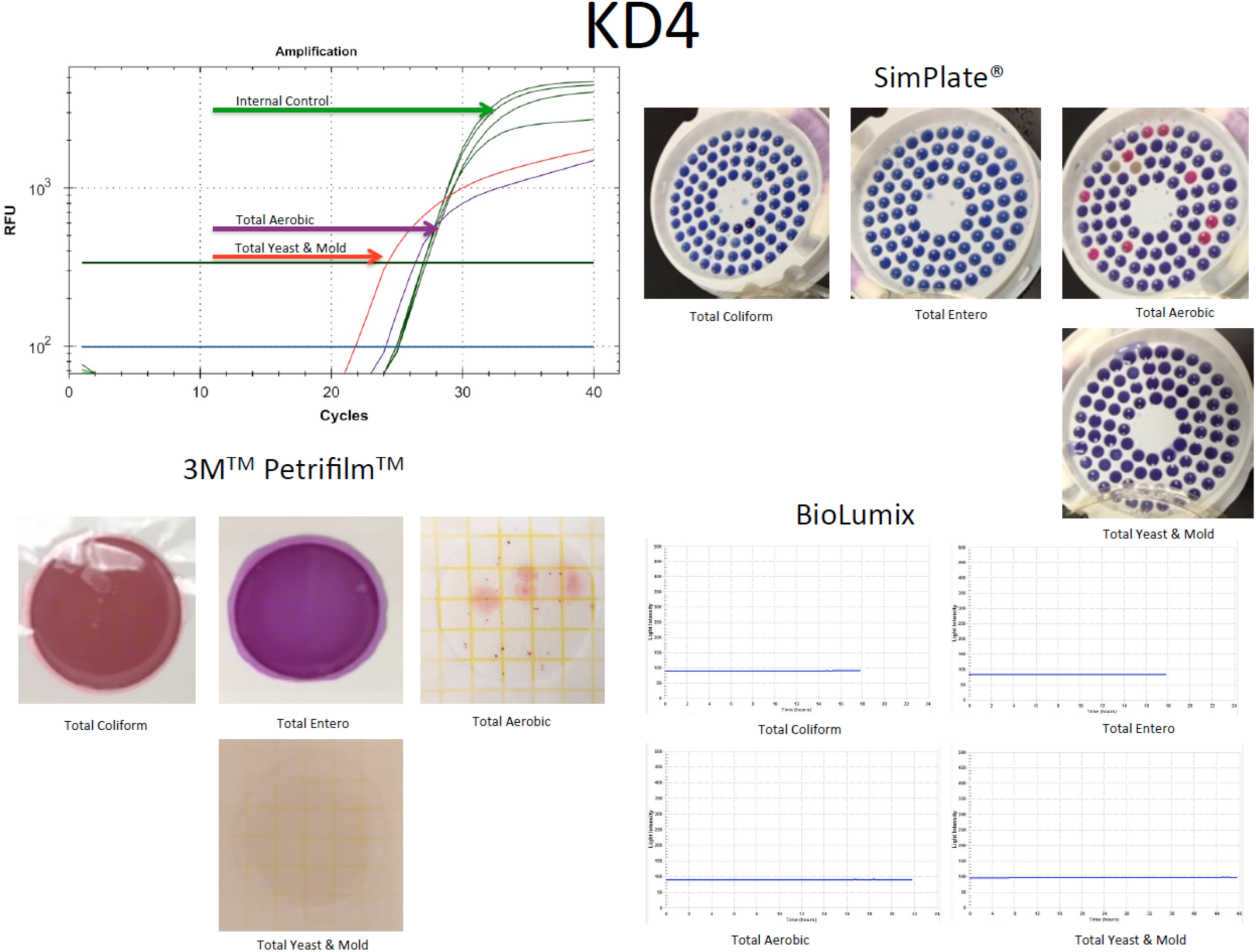
qPCR signal from TYM (red line) test run concurrently (multiplexed) with a plant internal control marker (green line). This marker targets a conserved region in the cannabis genome and should show up in every assay (Upper left). SimPlates count the number of discolored wells (Purple to pink) as a proxy for CFU/gram. Only total aerobic show growth (Upper Right). Petrifilm only demonstrate colonies on Total Aerobic platings (lower left). Biolumix demonstrate no signal across all 4 test (lower right).

**Figure 1B.**
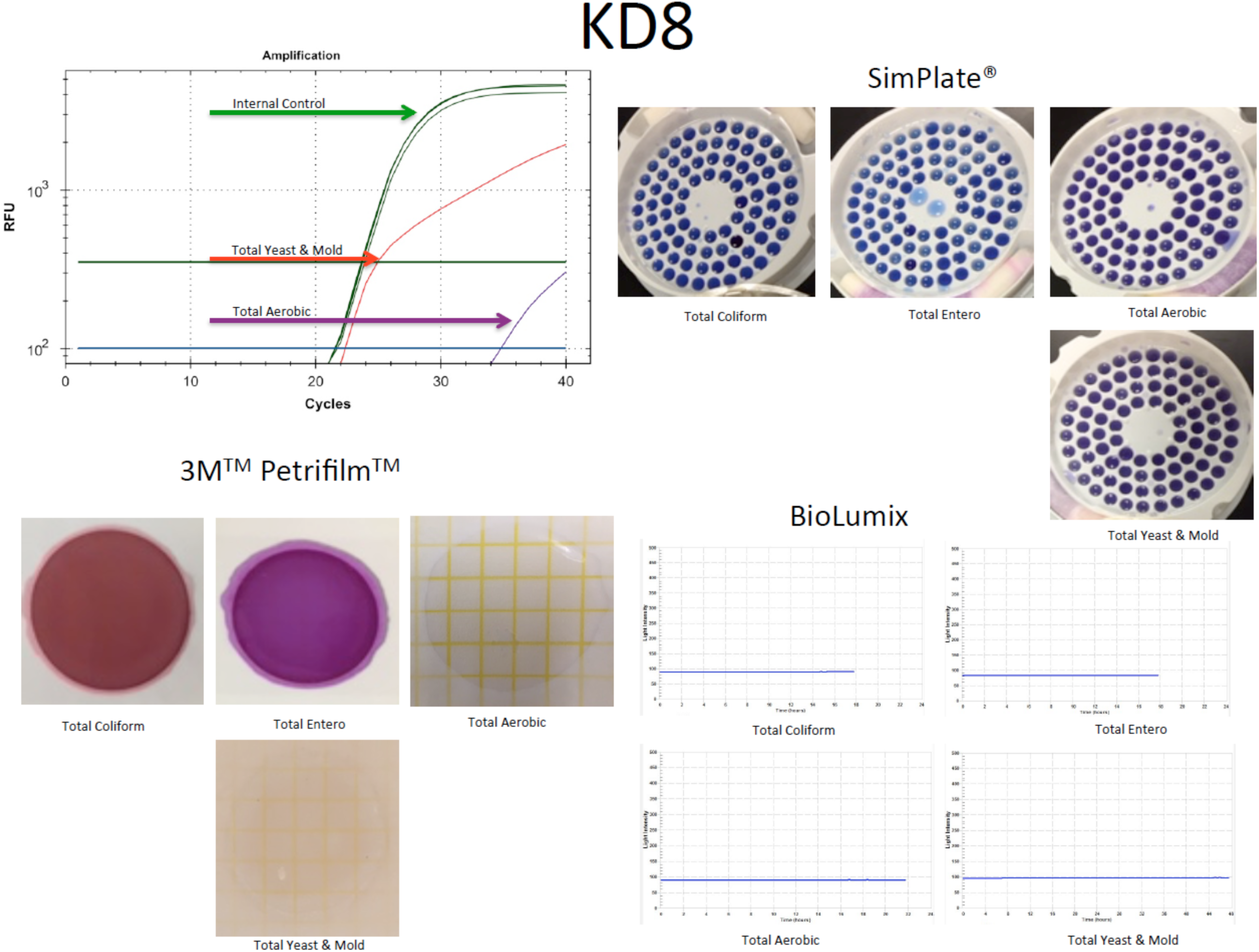
Sample KD8 fails to culture any Total Yeast and mold yet demonstrates significant TYM qPCR signal. Sample was graduated to ITS based next generation sequencing.

**Figure 1C.**
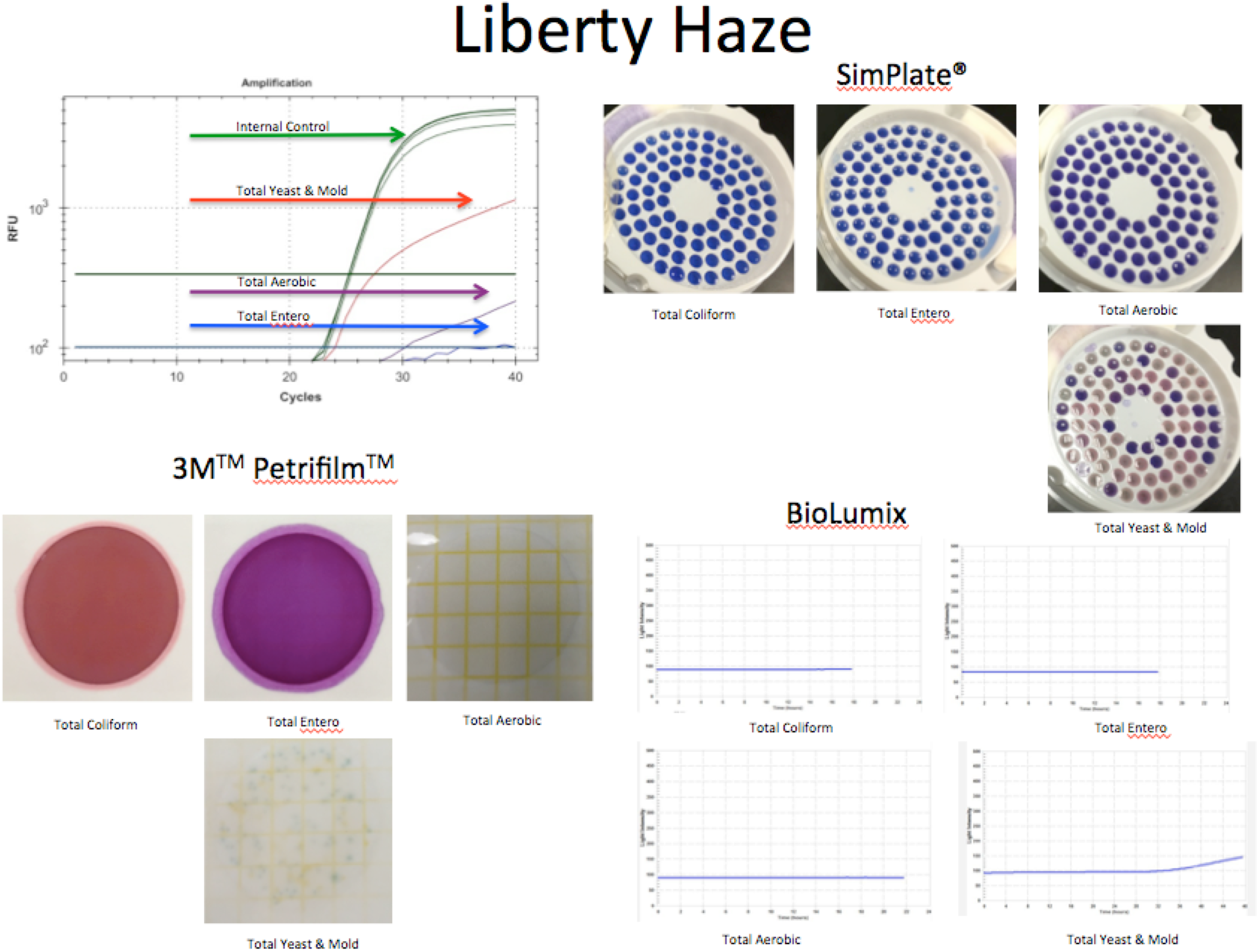
Sample Liberty Haze was tested with 3 culture based methods and compared to qPCR. Sample was graduated to ITS based next generation sequencing.

**Figure 2.**
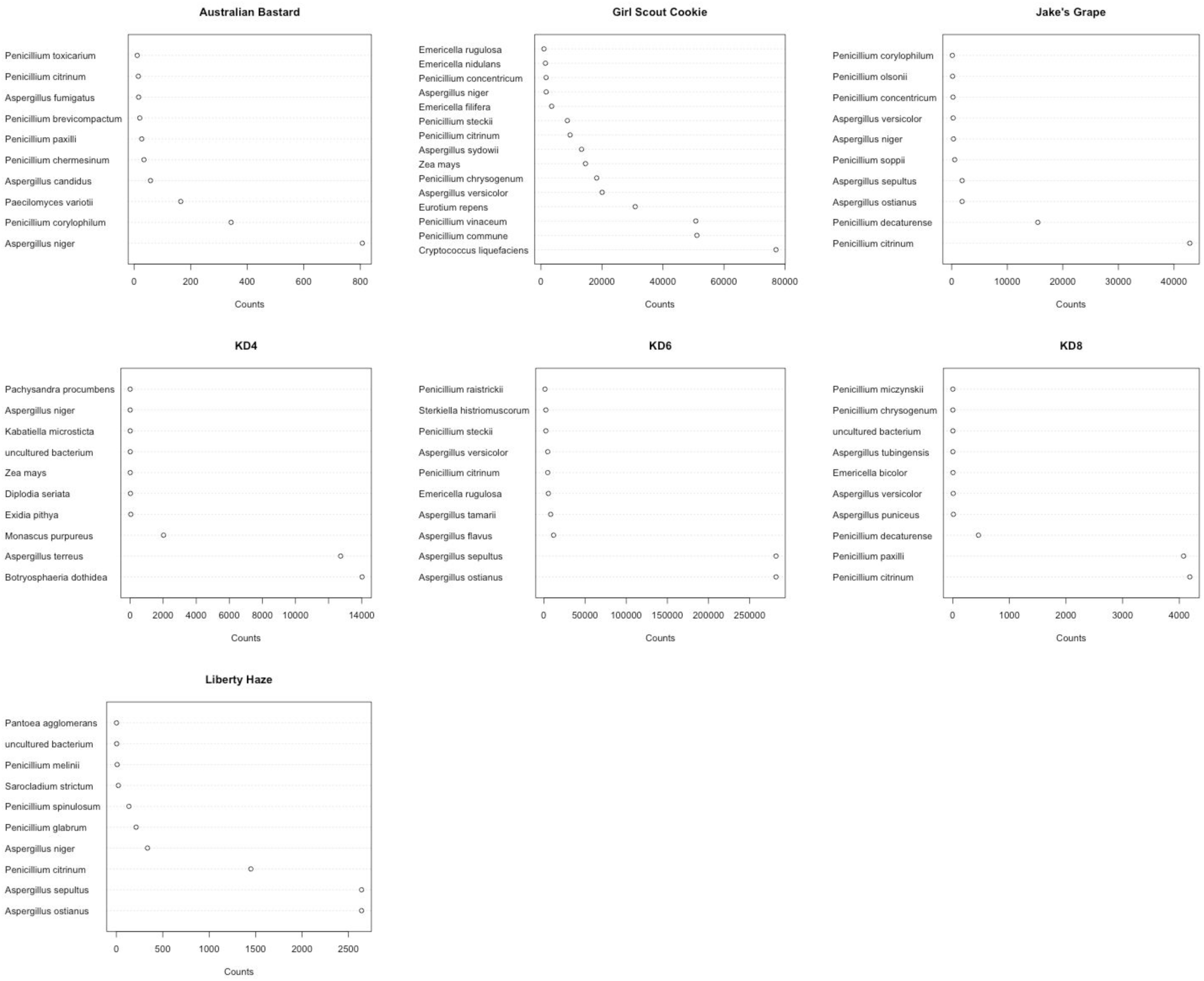
DNA sequencing of ITS3 amplicons from culture negative samples that are qPCR positive for Total Yeast and Mold tests. Penicillium and Aspergillus are commonly found (Y axis) but at different read counts in each sample (X axis). Read counts are more a reflection of sample normalization for sequencing than inter sample quantitation provided by qPCR.

We further analyzed the ITS sequence alignments using the whole genome shotgun based microbiome classification software known as One Codex ^33^. Nine of the ten samples sequenced showed the presence of *P. paxilli* (Figure 3). To verify the accuracy of this ITS phylotyping, a gene involved in the Paxilline toxin Biosynthesis pathway of *P. paxilli* was amplified with PaxPss1 and PaxPss2 primers described by Saikia *et al^34^.* The resulting 725bp amplicon (expected size) was sequenced to confirm the presence of the *P. Paxilli* biosynthesis gene in the cannabis sample KD8 (Figure 4). While there are some discrepancies between the two software platforms, our analysis used merged paired reads with MG-RAST and correlate better with PCR results. While One Codex predicted and confirmed KD8 as having the highest Paxilli content, the One Codex platform is optimized for whole genome shotgun data.

**Figure 3.**
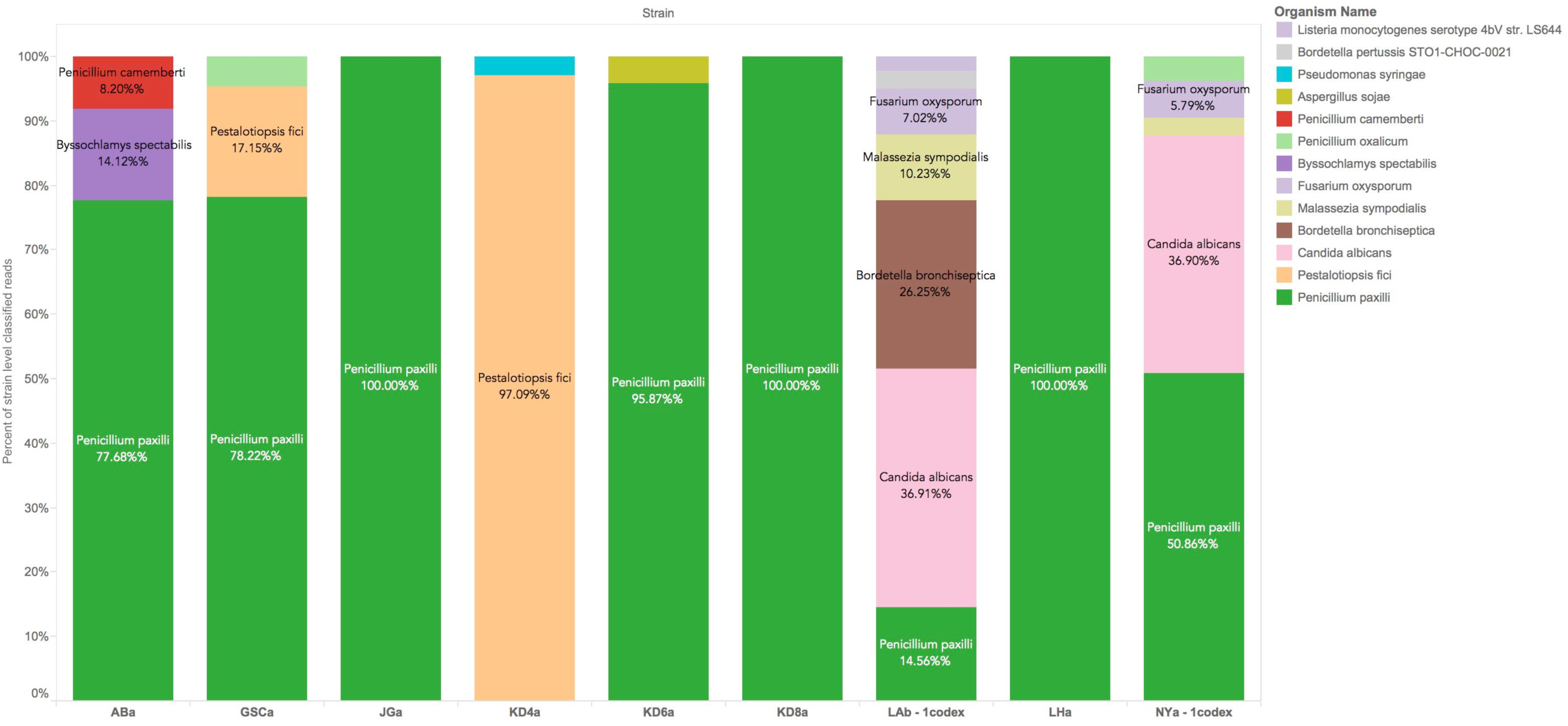
One Codex classification of ITS reads. P.Paxilli is the most frequently found contaminant in Cannabis flowers. P.Citrinum is not in the One Codex database at this time. One Codex utilizes a fast k-mer based approach for whole genome shotgun classification and can be influenced by read trimming and database content. The reads provided to MG-RAST were trimmed and FLASH’d (Paired end reads merged when overlapping) prior to classification. K-mer based approaches can significantly differ from longer word size methods and this underscores the importance of confirmatory PCR in microbiome analysis.

**Figure 4.**
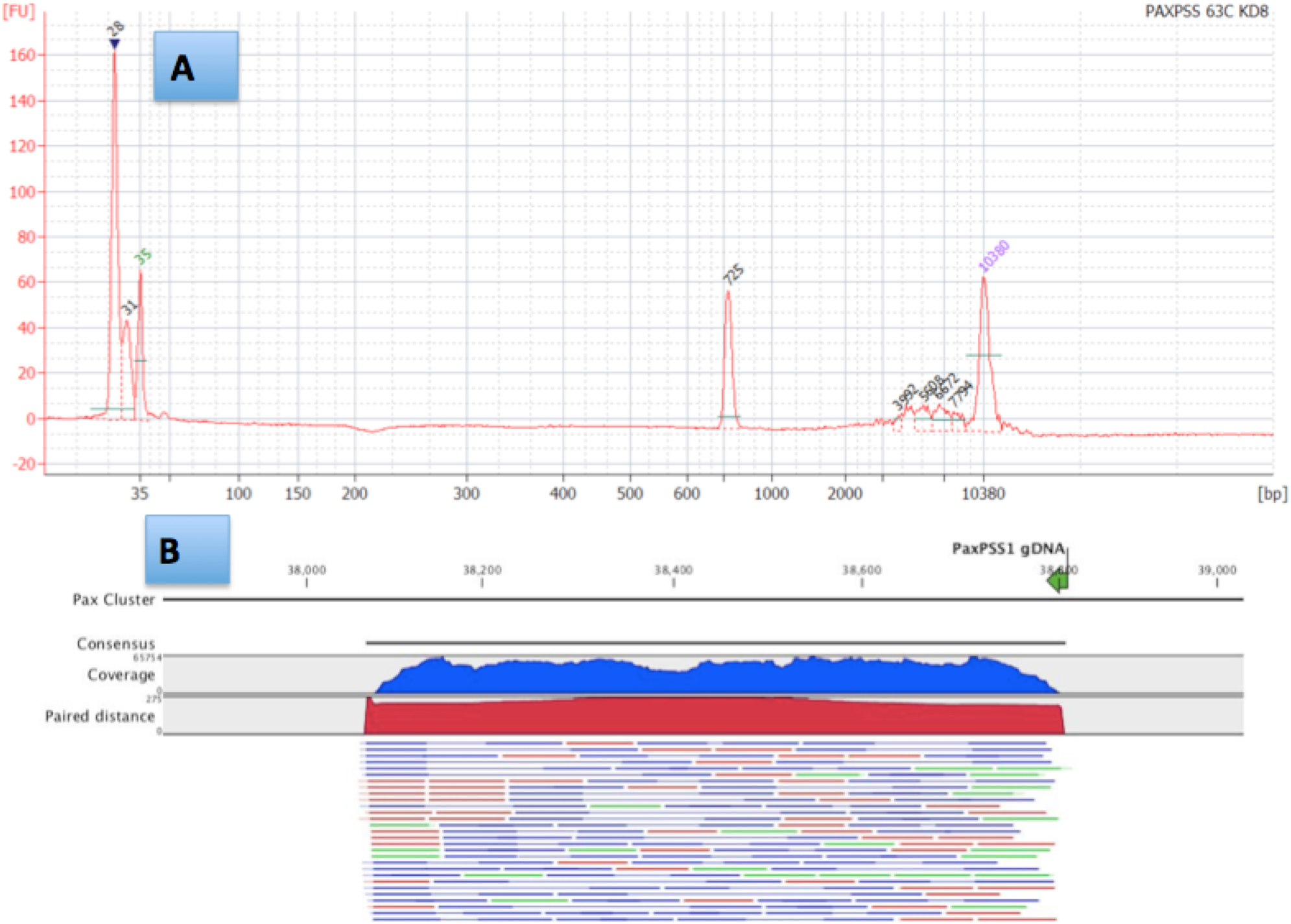
PaxP PCR demonstrates amplification of a 725bp band in sample KD8(A). PCR products were made into a shotgun library with Nextera and sequenced on an Illumina MiSeq with 2×75bp reads to over 10,000X coverage. Reads were mapped with CLCbio 4 to NCBI Accession number HM171111.1. Paired reads are displayed as blue lines, Green and Red lines are unpaired reads. Read Coverage over the amplicon is depicted in a blue histogram over the cluster while paired end read distance is measured in a red histogram over the region. Off target read mapping is limited (B). Alignment of PCR primers to P.Paxilli reference shows a 5 prime mismatch that is a result of the primers being designed to target spliced RNA according to Saikia *et al.*

With the confirmed presence of *P. paxilli*, we are curious to find out whether the toxin, paxilline, is present in the samples. Development of monoclonal antibodies to paxilline has recently been described^35^, but commercial ELISA assays with sensitivity under 50ppb do not appear to be available at this time. A >50ppb multiplexed ELISA assay is available from Randox Food Diagostics (Crumlin,UK). Detection with LC-MS/MS has also been described^36, 37^, however, and experiments are underway to determine whether Paxilline can be identified in the background of cannabinoids and terpenes present in Cannabis samples.

## Discussion

Several potentially harmful fungal species were detected in dispensary-derived Cannabis samples by qPCR and subsequent sequencing in this study. Three different culture-based assays failed to detect all of the positive samples and one, BioLumix^TM^, detected only one out of 7 positive samples. A review of the literature suggests that Penicillium microbes can be cultured on CYA media, but some may require colder temperatures (21-24C) and 7 day growth times^38^. Of the Penicillium, only P. Citrinum has been previously reported to culture with 3M Petri-Film^39^. In addition, several studies have demonstrated plant phytochemicals and terpenoids like eugenol can inhibit the growth of fungi ^40^. It is possible the different water activity of the culture assay compared to the natural terpene rich flower environment is contributing to the false negative test results.

Quantitative PCR is agnostic to water activity and can be performed in hours instead of days. The specificity and sensitivity provides important information on samples that present risks invisible to culture based systems. The draw back to qPCR is the method’s indifference to living or non-living DNA. While techniques exist to perform live-dead qPCR, the live status of the microbes is unrelated to toxin potentially produced while the microbes were alive. ELISA assays exists to screen for some toxins^41^. Current State-recommended ELISA’s do not detect Citrinin or Paxilline, the toxins produced by *P. citrinum* and *P. paxilli*, respectively. The predominance of these Penicillium species in a majority of the samples tested is interesting. Several Penicillium species are known to be endophytes on various plant species, including *P. citrinum* ^10^, and this raises the question of whether they are also Cannabis endophytes.

Paxilline is a tremorgenic and ataxic potassium channel blocker and has been shown to attenuate the anti-seizure properties of cannabidiol in certain mouse models^42-44^. Paxilline is reported to have tremorgenic effects at nanomolar concentrations and is responsible for Ryegrass-staggers disease^45^. Cannabidiol is often used at micromolar concentrations for seizure reduction implying sub-percentage contamination of Paxilline could still be a concern. Citrinin is a mycotoxin that disrupts Ca2+ efflux in the mitochondrial permeability transition pore (mPTP)^46-53^. Ryan *et al.* demonstrated that cannabidiol affects this pathway suggesting a potential concern for CBD-mycotoxin interaction ^54^ Considering the hydrophobicity of Paxilline and the recent interest in the use of cannabidiol derived from cannabis flower oils for drug resistant Epilepsy, more precise molecular screening of fungal toxins may be warranted ^55-60^.

Our survey of cannabis flowers in this study was limited. Further screening will be required to define a set of tests that can adequately capture all risks. While ELISA assays are easy point of use tests that can be used to detect fungal toxins, they can suffer from lack of sensitivity and cross reactivity. ITS amplification and sequencing offers hypothesis-free testing that can complement the lack of specificity in ELISA assays. Appropriate primer design can survey a broad spectrum of microbial genomes while affording rapid iteration of design. Quantitative PCR has also demonstrated single molecule sensitivity and linear dynamic range over 5 orders of magnitude offering a very robust approach for detection of microbial risks. This may be important for the detection of nanomolar potency mycotoxins. Further studies are required to validate better detection methods for these toxins and verify whether Paxilline or Citrinin are present on cannabis at concentrations that present a clinical risk.

## Conclusions

These results demonstrate that culture based techniques superimposed from the food industry should be re-evaluated based on the known microbiome of actual Cannabis flowers in circulation at dispensaries. Several mycotoxin producing molds were detected that can potentially interfere with the medical use of cannabidiol. These microbes failed to grow on traditional culture based platforms but were rapidly detected with molecular based techniques. Further studies are required to quantitate the presence and concentration of mycotoxin production.

## Methods

### Culture based methods

3.55ml of Tryptic Soy Broth (TSB) was used to wet 250mg of homogenized flower in a whirlpack bag. TSB was aspirated from the reverse side of the 100um mesh filter and placed into a Biolumix ^TM^ growth vial and spread onto a 3M Petri Film ^TM^ and a SimPlate ^TM^ according to the respective manufacturers recommendations. Biolumix ^TM^ vials were grown and monitored for 48 hours while Petri-films ^TM^ and SimPlates ^TM^ were grown for 5 days. Petri-films ^TM^ and SimPlates ^TM^ were colony counted manually by three independent observers. Samples were tested on Total Coliform, Total Entero, Total Aerobic, and Total Yeast and Mold. Only Total Yeast and Mold discrepancies were graduated to sequencing.

### DNA Purification

Plant DNA was extracted with SenSATIVAx according to manufacturers instructions (Medicinal Genomics part #420001). DNA is eluted with 50ul ddH20.

### Primers used for PCR and sequencing

PCR was performed using 5ul of DNA (3ng/ul) 12.5ul 2X LongAmp (NEB) with 1.25ul of each 10uM MGC-ITS3 and MGC-ITS3 primer (MGC-ITS3; TACACGACGTTGTAAAACGACGCATCGATGAAGAACGCAGC) and (MGC-ITS3R; AGGATAACAATTTCACACAGGATTTGAGCTCTTGCCGCTTCA) with 10ul ddH20 for a 25ul total reaction. An initial 95C 5 minute denaturization was performed followed by 40 cycles of 95C for 15s and 65C for 90s. Samples were purified with 75ul SenSATIVAx, washed twice with 100ul 70% EtOH and bench dried for 5minutes at room temperature. Samples were eluted in 25ul ddH20.

### Tailed PCR Cloning and Sequencing

DNA libraries were constructed with 250ng DNA using NEB’s NEBNext Quick ligation module (NEB # E6056S). End Repair used 3ul of Enzyme Mix, 6.5ul of Reagent Mix, 55.5ul of DNA + ddH20. Reaction was incubated at 30C for 20 minutes. After End Repair, Ligation was performed directly with 15ul of Blunt End TA Mix, 2.5ul of Ilumina Adaptor (10uM) and 1ul of Ligation enhancer (assumed to be 20% PEG 6000). After 15-minute ligation at 25C, 3ul of USER enzyme was added to digest the hairpin adaptors and prepare for PCR. The USER enzyme was tip-mixed and incubated at 37C for 20 minutes. After USER digestion, 86.5ul of SenSATIVAx was added and mixed. The samples were placed on a magnet for 15 minutes until the beads cleared and the supernatant could be removed. Beads were washed twice with 150ul of 70% EtOH. Beads were left for 10 minute to air dry and then eluted in 25ul of 10mM Tris-HCl.

### Library PCR

25ul 2X Q5 Polymerase was added to 23ul of DNA with 1ul of i7 index primer (25uM) and 1ul Universal primer (25um). After an initial 95C for 10 secs, the library was amplified for 15 cycles of 95C 10sec, 65C 90sec. Samples were purified by mixing 75ul of SenSATIVAx into the PCR reaction. The samples were placed on a magnet for 15 minutes until the beads cleared and the supernatant could be removed. Beads were washed twice with 150ul of 70% EtOH. Beads were left for 10 minute to air dry and then eluted in 25ul of 10mM Tris-HCl. Samples were prepared for sequencing on the MiSeq V2 chemistry according to the manufactures instructions. 2x250bp reads were selected to obtain maximal ITS sequence information.

### PaxP Verification PCR

Primers described by Shirazi-zand et al. were utilized to amplify a segment of the 725bp PaxP gene. 25ul LongAmp (NEB) 4ul 10uM Primer, 1ul DNA (14ng/ul), 20ul ddH20 to make a 50ul PCR reaction. Cycling conditions we slightly modified to accommodate a different polymerase. 95C for 30s followed by 28 cycles of 95C 15s, 55C for 30sec, 65C 2.5 minutes. Samples were purified with 50ul of SenSATIVAx as described above. 1ul of purified PCR product was sized on Agilent HS 2000 chip. Nextera libraries and sequencing were performed according to instructions from Illumina using 2x75bp sequencing on a version 2 MiSeq.

### Analysis

Reads were demultiplex and trimmed with Casava 1.8.2 and trim_galore. FLASH ^61^ was used to merge the reads using max_overlap 150. The reads were aligned to microbial references using MG-RAST^62^. Alignments and classifications were confirmed with a second software tool from One Codex and critical pathways identified for further evaluation with PCR of toxin producing genes. Reads are deposited in NCBI under SRA accession: SRP065410. Nextera 2x75bp sequencing of the PaxP gene was mapped to accession number HM171111.1 with CLCbio Workstation V4 at 98% identity over 80% of the read. One Codex analysis was put into Public mode under the following public URLs:

Australian Bastard:

https://app.onecodex.com/analysis/public/201e7f1642e04a3c

https://app.onecodex.com/analysis/public/58f1e03c10434bfa

KD4:

https://app.onecodex.com/analysis/public/2e86e262817246c4

https://app.onecodex.com/analysis/public/1abd5b60446140a0

KD6:

https://app.onecodex.com/analysis/public/a92d3dff5485499d

https://app.onecodex.com/analysis/public/8d72e2514e564ecd

KD8:

https://app.onecodex.com/analysis/public/8d72e2514e564ecd

https://app.onecodex.com/analysis/public/d6e2e0bcfba3469f

Liberty Haze:

https://app.onecodex.com/analysis/public/7bcd650fa5544f2c

https://app.onecodex.com/analysis/public/7f0feb6cb0a94d56

Girls Scout Cookie:

https://app.onecodex.com/analysis/public/a71b1ce8331c461d

https://app.onecodex.com/analysis/public/8d6f10c7ee684f93

Jakes Grape:

https://app.onecodex.com/analysis/public/bc8af5ed19e5407a

https://app.onecodex.com/analysis/public/99d7a4a2f7af486b

RECON:

https://app.onecodex.com/analysis/public/8a22a16cc2e24731

https://app.onecodex.com/analysis/public/0af6ae26a01f48d5

GreenCrack:

https://app.onecodex.com/analysis/public/6114843d2eb3425e

https://app.onecodex.com/analysis/public/3eee642786c54a88

LA Confidential:

https://app.onecodex.com/analysis/public/01e8aefb0d4f4f62

https://app.onecodex.com/analysis/public/b74c2988fcd84e38

NYC Diesel:

https://app.onecodex.com/analysis/public/441cfad759f64dcc

https://app.onecodex.com/analysis/public/d97b39cae96c4a44

## Author Contributions

KJM designed the study and performed the One-Codex analysis

JS designed and ran the culture and qPCR laboratory experiments

LZ assisted in the figure generation and laboratory experiment

YH assisted in sequencing and PCR confirmation of Pax.

VT-read alignment, MG-RAST, primer design and analysis.

TF-Sample tracking software, figure generations, ITS software comparisons

DS-Manuscript construction and review

## Acknowledgments

John McPartland, Cindy Orser, Brad Douglass, Joost Heeroma, Nick Greenfield and Kellie Dodd for thoughtful advice.

## References

1. Fujikawa H. & Itoh T. Tailing of thermal inactivation curve of Aspergillus niger spores. Applied and environmental microbiology 62, 3745–3749 (1996).

2. Kabak B. & Dobson A.D. Biological strategies to counteract the effects of mycotoxins. Journal of food protection 72, 2006–2016 (2009).

3. Rasooly R. & Do P.M. Shiga toxin Stx2 is heat-stable and not inactivated by pasteurization. International journal of food microbiology 136, 290–294 (2010).

4. Langouet S. et al. Metabolism of aflatoxin B1 by human hepatocytes in primary culture. Adv Exp Med Biol 387, 439–442 (1996).

5. Yamaori S., Ebisawa J., Okushima Y., Yamamoto I. & Watanabe K. Potent inhibition of human cytochrome P450 3A isoforms by cannabidiol: role of phenolic hydroxyl groups in the resorcinol moiety. Life Sci 88, 730–736 (2011).

6. Bragulat M.R., Martinez E., Castella G. & Cabanes F.J. Selective efficacy of culture media recommended for isolation and enumeration of Fusarium spp. Journal of food protection 67, 207–211 (2004).

7. Castella G., Bragulat M.R., Rubiales M.V. & Cabanes F.J. Malachite green agar, a new selective medium for Fusarium spp. Mycopathologia 137, 173–178 (1997).

8. Desjardins A.E. & Proctor R.H. Molecular biology of Fusarium mycotoxins. International journal of food microbiology 119, 47–50 (2007).

9. Stewart, E.J. Growing unculturable bacteria. Journal of bacteriology 194, 4151–4160 (2012).

10. Kusari P, K.S., Spiteller M, Kayser O Endophytic fungi harbored in Cannabis sativa L.: diversity and potential as biocontrol agents against host plantspecific phytopathogens. Fungal Diversity. 60, 137–151 (2013).

11. McPartland Fungal pathogens of Cannabis sativa in Illinois. Phytopathology 72:797 (1983).

12. McPartland Common names for diseases of Cannabis sativa L. PlantDis 75, 226–227 (1991).

13. McPartland Microbiological contaminants of marijuana. J. Int Hemp Assoc 1, 41–44 (1994).

14. McPartland Cannabis pathogens X: Phoma, Ascochyta and Didymella species. Mycologia 86, 870–878 (1995).

15. McPartland A review of Cannabis diseases. J Int Hemp Assoc 3, 19–23 (1996).

16. van Bakel H. et al. The draft genome and transcriptome of Cannabis sativa. Genome biology 12, R102 (2011).

17. McKernan The Cannabis Sativa Genome [Online] Available at: (https://aws.amazon.com/datasets/the-cannabis-sativa-genome/) Accessed on 13 September 2015. (2011).

18. McKernan [Online] Available at: http://www.icrs.co/SYMPOSIUM.2015/ICRS2015.Preliminary.Programme.pdf, Accessed on 20 October 2015 (2015).

19. Borneman, J. & Hartin, R.J. PCR primers that amplify fungal rRNA genes from environmental samples. Applied and environmental microbiology 66, 4356–4360 (2000).

20. White T.J., T. Bruns S. Lee & J. Taylor. Amplification and direct sequencing of fungal ribosomal RNA genes for phylogenetics. PCR Protocols: A Guide to Methods and Applications, 315–322 (1990).

21. Mason M., Cheung I. & Walker L. Creating a geospatial database of risks and resources to explore urban adolescent substance use. Journal of prevention & intervention in the community 37, 21–34 (2009).

22. Abd Alla E.A., Metwally M.M., Mehriz A.M. & Abu Sree Y.H. Sterigmatocystin: incidence, fate and production by Aspergillus versicolor in Ras cheese. Die Nahrung 40, 310–313 (1996).

23. Aly S.A. & Elewa N.A. The effect of Egyptian honeybee propolis on the growth of Aspergillus versicolor and sterigmatocystin biosynthesis in Ras cheese. The Journal of dairy research 74, 74–78 (2007).

24. Engelhart S. et al. Occurrence of toxigenic Aspergillus versicolor isolates and sterigmatocystin in carpet dust from damp indoor environments. Applied and environmental microbiology 68, 3886–3890 (2002).

25. Kocic-Tanackov S. et al. Effects of onion (Allium cepa L.) and garlic (Allium sativum L.) essential oils on the Aspergillus versicolor growth and sterigmatocystin production. Journal of food science 77, M278–284 (2012).

26. Song, F. et al. Three new sterigmatocystin analogues from marine-derived fungus Aspergillus versicolor MF359. Applied microbiology and biotechnology 98, 3753–3758 (2014).

27. El-Sayed Abdalla, A., Zeinab Kheiralla, M., Sahab, A. & Hathout A. Aspergillus terreus and its toxic metabolites as a food contaminant in some Egyptian Bakery products and grains. Mycotoxin research 14, 83–91 (1998).

28. Ames D.D., Wyatt R.D., Marks H.L. & Washburn K.W. Effect of citrinin, a mycotoxin produced by Penicillium citrinum, on laying hens and young broiler chicks. Poultry science 55, 1294–1301 (1976).

29. Mazumder P.M., Mazumder R., Mazumder A. & Sasmal D.S. Antimicrobial activity of the mycotoxin citrinin obtained from the fungus penicillium citrinum. Ancient science of life 21, 191–197 (2002).

30. Park S.Y. et al. Citrinin, a mycotoxin from Penicillium citrinum, plays a role in inducing motility of Paenibacillus polymyxa. FEMS microbiology ecology 65, 229–237 (2008).

31. Itoh Y., Johnson R. & Scott B. Integrative transformation of the mycotoxin-producing fungus, Penicillium paxilli. Current genetics 25, 508–513 (1994).

32. Shibayama M., Ooi K., Johnson R., Scott B. & Itoh Y. Suppression of tandem-multimer formation during genetic transformation of the mycotoxin-producing fungus Penicillium paxilli by disrupting an orthologue of Aspergillus nidulans uvsC. Current genetics 42, 59–65 (2002).

33. Minot S.S., Krumm N. & Greenfield N.B. One Codex: A Sensitive and Accurate Data Platform for Genomic Microbial Identification. bioRxiv (2015).

34. Saikia S., Parker E.J., Koulman A. & Scott B. Defining paxilline biosynthesis in Penicillium paxilli: functional characterization of two cytochrome P450 monooxygenases. The Journal of biological chemistry 282, 16829–16837 (2007).

35. Maragos C.M. Development and Evaluation of Monoclonal Antibodies for Paxilline. Toxins 7, 3903–3915 (2015).

36. Vishwanath V., Sulyok M., Labuda R., Bicker W. & Krska R. Simultaneous determination of 186 fungal and bacterial metabolites in indoor matrices by liquid chromatography/tandem mass spectrometry. Analytical and bioanalytical chemistry 395, 1355–1372 (2009).

37. Uhlig S., Egge-Jacobsen W., Vralstad T. & Miles C.O. Indole-diterpenoid profiles of Claviceps paspali and Claviceps purpurea from high-resolution Fourier transform Orbitrap mass spectrometry. Rapid communications in mass spectrometry: RCM 28, 1621–1634 (2014).

38. Houbraken J., Frisvad J.C. & Samson R.A. Taxonomy of Penicillium section Citrina. Studies in mycology 70, 53–138 (2011).

39. 3M http://multimedia.3m.com/mws/media/898592O/3m-petrifilm-rapid-yeast-mold-count-plate.pdf.

40. Zare M., Shams-Ghahfarokhi M., Ranjbar-Bahadori S., Allameh A. & Razzaghi-Abyaneh M. Comparative study of the major Iranian cereal cultivars and some selected spices in relation to support Aspergillus parasiticus growth and aflatoxin production. Iranian biomedical journal 12, 229–236 (2008).

41. Labs R. [Online] Available at: http://www.romerlabs.com/en/knowledge/mycotoxins/, Accessed on Oct, 2015.

42. Shirazi-zand Z., Ahmad-Molaei L., Motamedi F. & Naderi N. The role of potassium BK channels in anticonvulsant effect of cannabidiol in pentylenetetrazole and maximal electroshock models of seizure in mice. Epilepsy & behavior: E&B 28, 1–7 (2013).

43. Sabater-Vilar M., Nijmeijer S. & Fink-Gremmels J. Genotoxicity assessment of five tremorgenic mycotoxins (fumitremorgen B, paxilline, penitrem A, verruculogen, and verrucosidin) produced by molds isolated from fermented meats. Journal of food protection 66, 2123–2129 (2003).

44. Sanchez-Pastor E. et al. Cannabinoid receptor type 1 activation by arachidonylcyclopropylamide in rat aortic rings causes vasorelaxation involving calcium-activated potassium channel subunit alpha-1 and calcium channel, voltage-dependent, L type, alpha 1C subunit. European journal of pharmacology 729, 100–106 (2014).

45. Imlach W.L. et al. The molecular mechanism of “ryegrass staggers,” a neurological disorder of K+ channels. The Journal of pharmacology and experimental therapeutics 327, 657–664 (2008).

46. Chagas G.M., Campello A.P. & Kluppel M.L. Mechanism of citrinin-induced dysfunction of mitochondria. I. Effects on respiration, enzyme activities and membrane potential of renal cortical mitochondria. Journal of applied toxicology: JAT 12, 123–129 (1992).

47. Chagas G.M., Oliveira B.M., Campello A.P. & Kluppel M.L. Mechanism of citrinin-induced dysfunction of mitochondria. II. Effect on respiration, enzyme activities, and membrane potential of liver mitochondria. Cell biochemistry and function 10, 209–216 (1992).

48. Chagas G.M., Oliveira M.A., Campello A.P. & Kluppel M.L. Mechanism of citrinin-induced dysfunction of mitochondria. IV-Effect on Ca2+ transport. Cell biochemistry and function 13, 53–59 (1995).

49. Chagas G.M., Oliveira M.B., Campello A.P. & Kluppel M.L. Mechanism of citrinin-induced dysfunction of mitochondria. III. Effects on renal cortical and liver mitochondrial swelling. Journal of applied toxicology: JAT 15, 91–95 (1995).

50. Da Lozzo, E.J., Oliveira, M.B. & Carnieri E.G. Citrinin-induced mitochondrial permeability transition. Journal of biochemical and molecular toxicology 12, 291–297 (1998).

51. Jeswal, P. Citrinin-induced chromosomal abnormalities in the bone marrow cells of Mus musculus. Cytobios 86, 29–33 (1996).

52. Ribeiro S.M., Campello A.P., Chagas G.M. & Kluppel M.L. Mechanism of citrinin-induced dysfunction of mitochondria. VI. Effect on iron-induced lipid peroxidation of rat liver mitochondria and microsomes. Cell biochemistry and function 16, 15–20 (1998).

53. Ribeiro S.M., Chagas G.M., Campello A.P. & Kluppel M.L. Mechanism of citrinin-induced dysfunction of mitochondria. V. Effect on the homeostasis of the reactive oxygen species. Cell biochemistry and function 15, 203–209 (1997).

54. Ryan, D., Drysdale, A.J., Lafourcade, C., Pertwee, R.G. & Platt B. Cannabidiol targets mitochondria to regulate intracellular Ca2+ levels. The Journal of neuroscience: the official journal of the Society for Neuroscience 29, 2053–450 2063 (2009).

55. Devinsky O., Whalley B.J. & Di Marzo V. Erratum to: Cannabinoids in the Treatment of Neurological Disorders. Neurotherapeutics: the journal of the American Society for Experimental NeuroTherapeutics 12, 910 (2015).

56. Devinsky O., Whalley B.J. & Di Marzo, V. Cannabinoids in the Treatment of Neurological Disorders. Neurotherapeutics: the journal of the American Society for Experimental Neuro Therapeutics 12, 689–691 (2015).

57. Friedman, D. & Devinsky, O. Cannabinoids in the Treatment of Epilepsy. The New England journal of medicine 373, 1048–1058 (2015).

58. Rosenberg, E.C., Tsien, R.W., Whalley, B.J. & Devinsky, O. Cannabinoids and Epilepsy. Neurotherapeutics: the journal of the American Society for Experimental Neuro Therapeutics 12, 747–768 (2015).

59. Cilio M.R., Thiele E.A. & Devinsky, O. The case for assessing cannabidiol in epilepsy. Epilepsia 55, 787–790 (2014).

60. Devinsky, O. et al. Cannabidiol: pharmacology and potential therapeutic role in epilepsy and other neuropsychiatric disorders. Epilepsia 55, 791–802 (2014).

61. Magoc, T. & Salzberg S.L. FLASH: fast length adjustment of short reads to improve genome assemblies. Bioinformatics 27, 2957–2963 (2011).

62. Glass E.M., Wilkening, J., Wilke, A., Antonopoulos, D. & Meyer, F. Using the metagenomics RAST server (MG-RAST) for analyzing shotgun metagenomes. Cold Spring Harbor protocols 2010, pdb prot5368 (2010).

